# Doubly exponential decay of the ovarian reserve and aging milestones

**DOI:** 10.64898/2026.11.21.768290

**Authors:** Sean D. Lawley, Joshua Johnson

## Abstract

**Study question:** Is there a simple, quantitatively accurate, and mechanistically plausible framework for understanding the decay of the ovarian reserve and aging milestones across populations of women?

**Summary answer:** A “doubly exponential” model of ovarian reserve decay involving a simple mathematical formula closely fits histological follicle counts, recapitulates the population distributions of the age at natural menopause and the onset of the menopause transition, and may explain curious features of menopause timing.

**What is known already:** Quantitative descriptions of the decay of the ovarian reserve date back decades, but existing models require several fitted parameters, are cumbersome to apply or require computer simulation rather than a usable formula, lack plausible biological mechanisms, and/or do not address population variability in reserve size and menopause timing.

**Study design, size, duration:** This is a mathematical modeling study.

**Participants/materials, setting, methods:** This modeling study assembled a new dataset of histological follicle counts from 299 women in seven published studies and used published population distributions of the age at natural menopause and menopause transition, as well as published estimates of how a unilateral oophorectomy affects menopause timing.

**Main results and the role of chance:** Despite its simplicity, the new doubly exponential model provides a close fit to histological ovarian reserve data. Furthermore, the model recapitulates the observed population distributions for both the age at natural menopause and the preceding menopause transition, as well as much of the population variability observed in histological ovarian reserve data.

**Large scale data:** The follicle count dataset is provided as Supplementary Data.

**Limitations, reasons for caution:** Follicle count data were cross-sectional across women rather than longitudinal within individual women, and these data were pooled across studies using different histological counting protocols.

**Wider implications of the findings:** The model may explain some curious features of menopause timing, including the relatively narrow menopause age distribution despite wide variability in reserve size at birth across women and the surprisingly modest hastening of menopause induced by a unilateral oophorectomy. Doubly exponential decay arises if individual primordial follicles exit the reserve at independent, Gompertz-distributed random times, which is consistent with stochastic threshold-crossing mechanisms of follicle loss. The model may be used to quantitatively study emerging therapies that target ovarian aging.

## 1 Introduction

Human reproduction depends on a finite pool of oocytes called the ovarian reserve. Each oocyte is housed within a quiescent structure called a primordial follicle (PF). The ovarian reserve decays over a woman’s lifetime, from a “starting supply” of approximately *N* = 10^5^ to *N* = 10^6^ PFs at birth to only around *k* = 10^3^ PFs at menopause. The age at natural menopause (ANM; the age at the final menstrual period, identified retrospectively after 12 months of amenorrhea [39]) and the preceding age at menopause transition onset (AMT; age when menstrual cycles become irregular [19]) vary between women, and this variation is thought to depend jointly on the ovarian reserve starting supply and loss rate.

While the basic ovarian reserve loss pattern approximating exponential decay is well-recognized, the developmental and physiological processes that produce this pattern are not well understood. For instance, ovarian reserve size varies greatly between individuals. The population median at birth is *N* = 6.5*×* 10^5^ [20], but this starting supply varies by an order of magnitude between newborn girls. Furthermore, this variability appears to increase with age, as the ovarian reserve between age-matched women in their forties varies by around two orders of magnitude [18]. Additionally, in light of the aforementioned population variability in reserve size (i.e. roughly 1, 000% at birth and 10, 000% at midlife), the ANM population distribution is curiously narrow, as most women have ANM within around 20% of the population median of *T* = 51 years. The ANM is also remarkably robust to insults to the ovarian reserve, as removing an entire ovary (i.e. roughly 50% of the reserve) appears to hasten menopause by only around 2 years (i.e. less than 5%) [38].

In addition to outstanding questions of basic physiology, novel interventions targeting the ovarian reserve are being developed. Indeed, given the importance of maintaining the ovarian reserve for fertility, health span, and even lifespan [42], a variety of clinical interventions have been proposed to delay depletion of the reserve (see [21] for a review), including pharmaceutical, cell therapy, and surgical approaches. Pharmaceutical approaches use small molecules or biologic agents to slow PF activation by targeting stress-response, mitochondrial, or cell-cycle pathways. Cell therapy approaches aim to boost the reserve size using induced pluripotent or oogonial stem cells to generate new PFs. The surgical approach of ovarian tissue cryopreservation and autologous transplantation harvests and freezes a portion of the reserve at a younger age and reimplants it later in life to extend ovarian function [22].

A quantitative understanding of the decline of the ovarian reserve is essential to addressing the physiological and medical questions outlined above. Mathematical modeling of ovarian aging has a long history, dating back at least to the stochastic models of Faddy, Jones, and Edwards [13] and Faddy, Gosden, and Edwards [11]. These early models of mouse ovarian aging led to the influential “biphasic” model [12], which posited that the ovarian reserve in women decays exponentially at a constant rate until a woman’s late thirties, at which point the exponential decay rate abruptly accelerates. Since abrupt “step changes in physiological activity are rare” [10], Faddy and Gosden later posited a differential equation model in which the reserve decay rate gradually increases over time [10]. Later ovarian reserve size models with gradual changes in follicle loss rate include the compressed exponential model of Hansen et al. [18] and the Gaussian cumulative model of Wallace and Kelsey [40]. Several additional models were posited in Refs. [7, 20, 32, 34].

These prior models have fundamentally shaped our quantitative understanding of ovarian aging. Building on this foundation, we see room for further refinement regarding (i) the number of fitted parameters, (ii) the ovarian reserve data sets that the models were fit to, (iii) plausible biological mechanisms, (iv) complicated or intractable formulas, and (v) population variability in starting supply, reserve size later in life, AMT, and ANM.

For (i), not counting a parameter setting the starting supply *N* at birth, the biphasic model [12] and the differential equation model [10] each used three fitted parameters, the Gaussian cumulative model used four fitted parameters [40], and the compressed exponential model used two fitted parameters [18]. Fitted parameters are quantities in a model whose values are not measured or known in advance, but are instead tuned until the model matches the data. While different numbers of fitted parameters can be justifiable, fewer is generally preferable: a model with many adjustable parameters can fit almost any dataset, which makes the fit less meaningful and less likely to generalize. For (ii), while the Gaussian cumulative model was fit to 288 postnatal follicle counts (plus 37 prenatal counts), the biphasic, differential equation, and compressed exponential models were fit to significantly smaller data sets (110, 110, and 122 follicle counts, respectively [10, 12, 18]). For (iii), while the differential equation model was based on the physiological hypothesis that paracrine factors regulate the “ovarian economy,” the Gaussian cumulative model was employed because it involves a flexible mathematical equation that can be tuned to fit data sets from a variety of biophysical sources. For (iv), the differential equation model has no explicit formula for the ovarian reserve size as a function of time, and the Gaussian cumulative model involves products of complicated Gauss error functions. Hence, it is difficult to see how ovarian aging depends on the various parameters or the proposed clinical interventions. For (v), these prior models leave room for refinement regarding the aforementioned population variability, especially the relationships between variability in starting supply, variability in reserve size later in life, and variability in ANM and AMT.

In this paper, we first assemble a new dataset of histological follicle counts, where each count is a total per woman and is traced to a specific table or figure in a prior publication. The dataset includes 299 women from seven studies and is, to our knowledge, the largest compilation of follicle counts after birth [28]. We then propose a “doubly exponential” model of the decay of the ovarian reserve in women, which involves only a single fitted rate parameter in addition to the starting supply. We asked whether this model (a) fits follicle counts as closely as prior models while using fewer fitted parameters, (b) recapitulates the observed population distributions of the ANM and AMT when the starting supply and decay rate vary between women, and (c) accounts for the modest hastening of menopause after unilateral oophorectomy.

## 2 Materials and methods

### 2.1 Study design

This is a mathematical modeling study of previously published data. We defined the doubly exponential model of ovarian reserve decay, estimated its single rate parameter from histological follicle counts and compared its fit with prior models, added between-woman variability in the starting supply and the decay rate, and compared model predictions to published estimates of the population distributions of the ANM and AMT and how a unilateral oophorectomy affects the ANM. Table 1 details model parameters.

**Table 1:** Parameters of the doubly exponential model and the menopause models. The models and model parameters correspond to a pair of ovaries.

| Parameter | Symbol | Value | How the value was determined |
|---|---|---|---|
| Median starting supply | *N*<sub>median</sub> | 6.5 × 10<sup>5</sup> follicles | Fixed from a prior analysis [20] |
| Variability of starting supply | *σ*<sub>*N*</sub> | 0.5 | Fixed from a prior analysis [20] |
| Median decay rate | *R*<sub>median</sub> | 0.0396 year<sup>−1</sup> | Fitted to 292 per-woman follicle counts (this study) |
| Variability of decay | *σ*<sub>*R*</sub> | 0.05 | Calibrated to observed ANM distributions [33, 41] |
| ANM threshold (threshold model) | *k* | 1000 follicles | Fixed at the conventional value [10, 12, 18] |
| AMT threshold (threshold model) | *k*<sub>MT</sub> | 2600 follicles | Calibrated to the observed median AMT [35] |
| Gap length for ANM (gaps model) | *g*<sub>ANM</sub> | 7.25 days | Calibrated so median ANM agrees with the threshold model |
| Gap length for AMT (gaps model) | *g*<sub>MT</sub> | 3.75 days | Calibrated so median AMT agrees with the threshold model |

### 2.2 Data sources

#### Histological follicle counts

We fitted the model to histological follicle counts from 292 women. The highly cited compilation of Wallace and Kelsey [40] made historical counts widely accessible, and earlier analyses relied on this important compilation [20,26,27]. However, in tracing the data back to the original publications, we found some inconsistencies (see the Supplemental Information (SI)). We therefore compiled our own dataset from prior publications.

Each count in our dataset is the estimated total for one woman (i.e. the total across both ovaries) and is traced to a specific table or figure in a prior publication (Supplementary Data [28]). Our fitting set includes nonzero follicle counts obtained at or after birth, regardless of menstrual status, and excludes counts of germ cells rather than follicles [2, 3], group means, and counts from infants born preterm. The 292 counts in our fitting set came from the following seven studies: Block [4] (43 women) and Block [5] (7 newborn infants), Richardson et al. [37] (14 women), Gougeon et al. [16] (43 women, plus 9 women shown only in figures by Faddy and colleagues [9, 12]), Forabosco and Sforza [14] (6 infants), Hansen et al. [18] (118 women and infants), and Knowlton et al. [25] (52 women). Seven women with zero follicles (3 in [37] and 4 in [18]) were excluded because they cannot enter a fit on the logarithmic scale, but we examine their effect in the SI. The counts of Knowlton et al. [25] and of the 9 women of Gougeon not tabulated by Gougeon et al. [16] were estimated from published figures [9, 12, 25] because their numerical counts were not published. The studies used somewhat different follicle definitions (for example, Block counted primordial follicles, whereas Hansen et al. [18] and Knowlton et al. [25] counted primordial, intermediate, and primary follicles). For the studies in the compilation of Wallace and Kelsey [40], we used their definition, and for Knowlton et al. [25], we used the same definition as Hansen et al. [18]. We examine alternative definitions in the SI.

#### Definition of the ovarian reserve: one versus two ovaries

In this paper, the “ovarian reserve” refers to the total number of follicles in a woman (i.e. the total across both ovaries), and thus *F* (*t*), *N*, *k*, and *k*_MT_ below refer to this total. This “per-woman” definition follows Refs. [14, 18], whereas Ref. [40] reported counts per ovary. In our dataset, both ovaries were counted and summed for 92 women [4, 5, 16]. For the 118 women of Hansen et al. [18], one or both ovaries were counted and the authors summed or doubled them. For 73 women, one ovary was counted and doubled, either by the original authors [5, 25] or by us [14, 37]. Faddy and colleagues plotted the remaining 9 counts per pair of ovaries [9, 12]. Doubling a count from a single ovary is unbiased if a woman’s two ovaries do not differ systematically, as found by Refs. [5, 15, 18]. For the 92 women whose two ovaries were counted separately [4, 5, 16], twice the count of one ovary was typically within 30% of the total, which is small compared to the roughly 3.5-fold scatter of the counts around the fitted curve. Finally, fitting counts per ovary with starting supply *N/*2 gives the identical rate constant.

#### Starting supply

The population median starting supply *N* = 6.5*×* 10^5^ and its population distribution (see “Population variability” below) were taken from a previously reported analysis of 30 counts taken from girls within a few months of birth [20].

#### Menopause timing

Empirical population distributions of the ANM used for comparison in Figure 2A were taken from Refs. [33, 41]. The empirical population distribution of the AMT used in Figure 2B was taken from Ref. [35]. Data on the effect of removing one ovary on menopause timing were taken from Ref. [38].

### 2.3 Doubly exponential model of ovarian reserve decay

The doubly exponential model posits that the median size of the ovarian reserve (total across both ovaries) in a population of women at age *t* ≥ 0 (time since birth) is

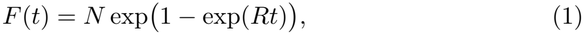

where *N* is the population median starting supply and *R >* 0 is a rate constant. We take *N* = 6.5 *×*10^5^, which is the previously observed population median starting supply [20], and thus the rate constant *R* is the only parameter fitted to follicle count data.

### 2.4 Model fitting and comparison

The rate constant *R* in Eq. (1) was estimated by minimizing the sum of squared errors (SSE) between *F* (*t*) and the follicle counts on a logarithmic scale (the SSE is computed with a base 10 logarithm). The starting supply was held at *N* = 6.5 *×*10^5^.

We compared the doubly exponential model with eight previously published models of ovarian reserve decay: the biphasic model [12], the ordinary differential equation (ODE) model [10], the compressed exponential model [18], the Gaussian cumulative model [40], the Gompertz model [7], the random walk model [20], and two chemical-kinetic models [32, 34]. The prior models are reviewed in the SI. Each prior model was evaluated on the same 292 counts, both at its original parameter values and after refitting its parameters to these counts. Refitting was done because the prior models were fit to smaller data sets (or a data set including prenatal counts [40]). Since the models have different numbers of fitted parameters, we followed Ref. [7] and compared them using the corrected Akaike information criterion, denoted AIC_c_ [1, 6], which penalizes additional fitted parameters. Further details are given in the SI.

### 2.5 Population variability

We interpret Eq. (1) with the median parameter values as describing the “median woman” in a population. To model a population of women, we allow the starting supply and the decay rate to vary between women. Based on a previously reported analysis of 30 follicle counts from girls within a few months of birth [20], we assume that the starting supply *N* varies across a population of women according to a log-normal distribution defined by the following cumulative distribution function,

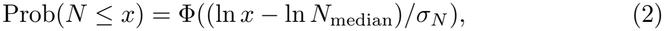

where ln is the natural logarithm, Φ is the cumulative distribution function of the standard normal distribution, and the median and variability parameter are *N*_median_ = 6.5 *×*10^5^ and *σ*_*N*_ = 0.5, respectively. The bottom 1% and top 1% of this distribution are *N*_1%_ *≈* 2 *×*10^5^ and *N*_99%_ *≈*2 *×*10^6^, respectively.

In some of our analysis, we take the rate constant to be the same for all women, *R* = 0.0396 year^*−*1^ (see Results). In the remaining analysis, since it is reasonable to suppose that the rate constant varies across the population, we assume that *R* varies according to a log-normal distribution defined by the following cumulative distribution function,

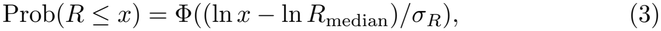

where the median is *R*_median_ = 0.0396 year^*−*1^ and the variability parameter is *σ*_*R*_ = 0.05. The coefficient of variation (the ratio of the standard deviation to the mean of a log-normally distributed random variable with variability parameter *σ* is 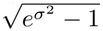 *≈ σ* for small *σ*, so *σ*_*R*_ = 0.05 means that most of the population is within 5% of the median value of *R*. The value *σ*_*R*_ = 0.05 was chosen so that the simulated ANM distribution resembles the observed distributions [33, 41] (see Results). For simplicity, we assume that a woman’s starting supply *N* is independent of her decay rate *R*.

### 2.6 Timing of menopause, the menopause transition, and the effect of unilateral oophorectomy

#### Threshold model

It has been assumed that the ANM corresponds to the time when the ovarian reserve size hits a critical threshold [10, 12, 18, 40]. Mathematically, this means that the ANM is the age *T* such that

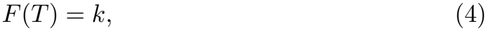

where *F* (*t*) is the ovarian reserve size at age *t* ≥ 0 and *k* is the critical threshold. Following [12], we set *k* = 1000 throughout this paper. The simple formula for *F* (*t*) in Eq. (1) means that it is straightforward to solve (4) for *T* to obtain

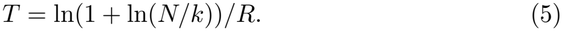

The formula (5) shows how the ANM *T* depends on the starting supply *N*, the rate constant *R*, and the threshold *k*. Analogously, we suppose that the menopause transition (MT) begins when the ovarian reserve hits a threshold *k*_MT_, so that the AMT (denoted *T*_MT_) is

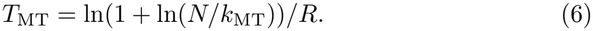

We set *k*_MT_ = 2600, which gives the observed median AMT (see Results). Some models assume that the ovarian reserve decays stochastically over time *t ≥* 0 with mean *F* (*t*), and thus the ANM *T* is defined as the stochastic time in which the reserve size is *k* [26, 31]. *However, we show in the SI that such a stochastic time has a mean that is very well approximated by the value of T* satisfying (4) and that, furthermore, the fluctuations about the mean are only a few months. Population distributions of *T* and *T*_MT_ were obtained by evaluating Eqs. (5)-(6) at values of *N* and *R* drawn independently from Eqs. (2)-(3).

#### Gaps model

An alternative “gaps” model of the timing of menopause and the MT was recently proposed [30] that offers some advantages when compared to the historical threshold assumption. In this model, menopause is triggered the first time that *g*_ANM_ consecutive days pass in which no follicle leaves the ovarian reserve. The physiological rationale for the dependence of MT and ANM timing upon stochastically appearing gaps of increasing length with chronological age is as follows. Since growing follicles are required to produce endocrine factors necessary for menstrual cycles, a sustained absence of newly growing follicles would disrupt cycling entirely; lengthy enough “gaps” thus trigger menopause. Similarly, the MT begins at the first gap of *g*_MT_ days, which is a shorter gap that disrupts cycle regularity [30].

We apply the gaps model to the entire ovarian reserve (i.e. both ovaries). In this model, each of the *N* follicles of a woman leaves the reserve at an independent random time *τ* with probability distribution Prob(*τ > t*) = exp(1*−* exp(*Rt*)), and thus the expected size of the reserve is *F* (*t*) in Eq. (1), and the ANM (respectively, AMT) is the first time that a time *g*_ANM_ elapses (respectively, *g*_MT_) in which no follicle leaves the reserve. Each simulated woman has a starting supply *N* and rate parameter *R* sampled independently from Eqs. (2)-(3), and the ANM and AMT distributions are computed from 10^5^ simulated women. We chose *g*_ANM_ = 7.25 days and *g*_MT_ = 3.75 days so that the median ANM and AMT of the gaps model equal those of the threshold model with *k* = 1000 and *k*_MT_ = 2600.

#### Unilateral oophorectomy

We model a unilateral oophorectomy (UO) as the instantaneous removal of half of the follicles in the reserve at the age of surgery. We assume that follicles in the remaining ovary continue to leave the reserve according to the same process as before surgery (i.e. there is no compensation by the remaining ovary) and that menopause occurs when the remaining reserve reaches the same threshold *k* as in women with two ovaries. These are simplifying assumptions since the remaining ovary may compensate for the removed ovary, perhaps through changes in follicle activation, recruitment, or endocrine feedback [24]. Furthermore, since a woman’s two ovaries might not contain equal numbers of follicles, we also predict how a UO affects menopause age if one ovary contains more or less than half of the reserve.

## 3 Results

### 3.1 One fitted rate describes follicle loss

Fitting the doubly exponential model in Eq. (1) to the 292 nonzero histological follicle counts after birth from seven studies (each a total for both ovaries) gives *R* = 0.0396 year^*−*1^. Figure 1 compares the resulting doubly exponential model to the 292 per-woman counts, together with the biphasic model [12], the ODE model [10], the compressed exponential model [18], and the Gaussian cumulative model [40].

**Figure 1:**
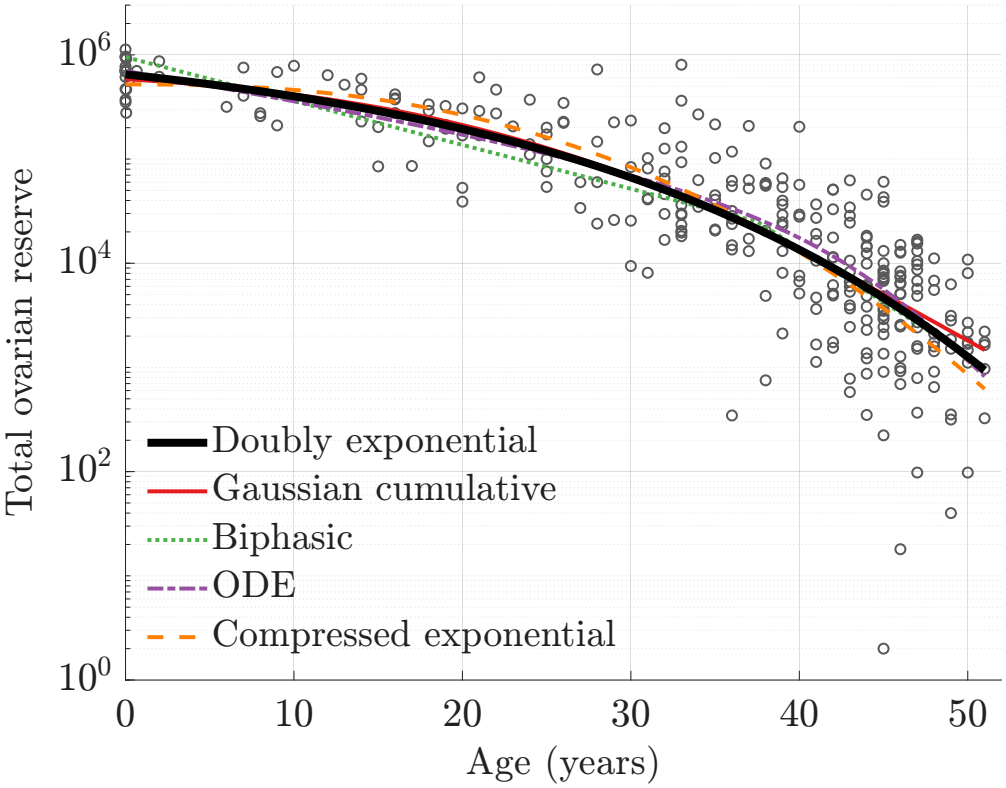
Comparison of the doubly exponential model (Eq. (1) with *R* = 0.0396 year^*−*1^) with the Gaussian cumulative model [40], biphasic model [12], ODE model [10], and compressed exponential model [18], each at its published parameter values. The circle markers are histological follicle counts for 292 women [4, 5, 9, 12, 14, 16, 18, 25, 37]. All counts and model curves are totals per woman (both ovaries). Seven counts of zero are not shown.

Using the parameter values reported in the original publications, the doubly exponential model has the best fit of all the models (we measure fit by the sum of squared errors between the model and the counts on a logarithmic scale). However, unlike the published parameters of the other models, the rate constant of the doubly exponential model was fitted to these counts. After refitting the other models to these counts, the four-parameter ODE model [10] attains the best fit, but the fit of the one-parameter doubly exponential model is only one half of one percent worse. Hence, the doubly exponential model attains a practically identical fit as the best fitting prior model while using only one fitted parameter rather than four fitted parameters. Following Ref. [7], this advantage can be quantified using the statistical estimator AIC_c_ [1, 6], which rewards a close fit but penalizes each additional fitted parameter. Indeed, the doubly exponential model has a better AIC_c_ than any prior model. Table S2 in the SI summarizes the comparison with prior models.

### 3.2 Menopause timing matches population data

Taking the population median values of *N* = 6.5 *×* 10^5^ and *R* = 0.0396 year^*−*1^ in the doubly exponential model implies that the ovarian reserve hits the conventional menopause threshold of *k* = 1000 follicles at age *T* = 50.8 years, which agrees with the empirically observed median ANM across a population of about 51 years [33, 41]. Similarly, the menopause transition occurs at age *T*_MT_ = 47.4 years if we take *k*_MT_ = 2600 in Eq. (6), which agrees with the empirically observed median AMT of 47.3 years [35].

If we allow the starting supply *N* to vary according to the empirically observed log-normal distribution in Eq. (2) but keep *R* fixed, then the median of the simulated ANM distribution agrees with the observed value of 51 years, but the distribution is less variable than what is observed in real populations [33,41]. The solid blue curve in Figure 2A is the distribution of the ANM when both *N* and *R* vary (Eqs. (2)-(3)). We make three comments about this result. First, by positing some population variability in *R* in addition to the empirically observed variability in *N*, the predicted ANM distribution resembles the distribution observed in actual populations (red plus markers [33] and red circle markers [41]). Second, the required variability in *R* is rather small. Indeed the value *σ*_*R*_ = 0.05 used in Figure 2A means that most of the population is within 5% of the median value of *R*, and essentially all of the population is within 3*σ*_*R*_ = 15% of the median. Third, other distributions of *R* centered at 0.0396 year^*−*1^ with small coefficients of variation yield similar ANM distributions.

**Figure 2:**
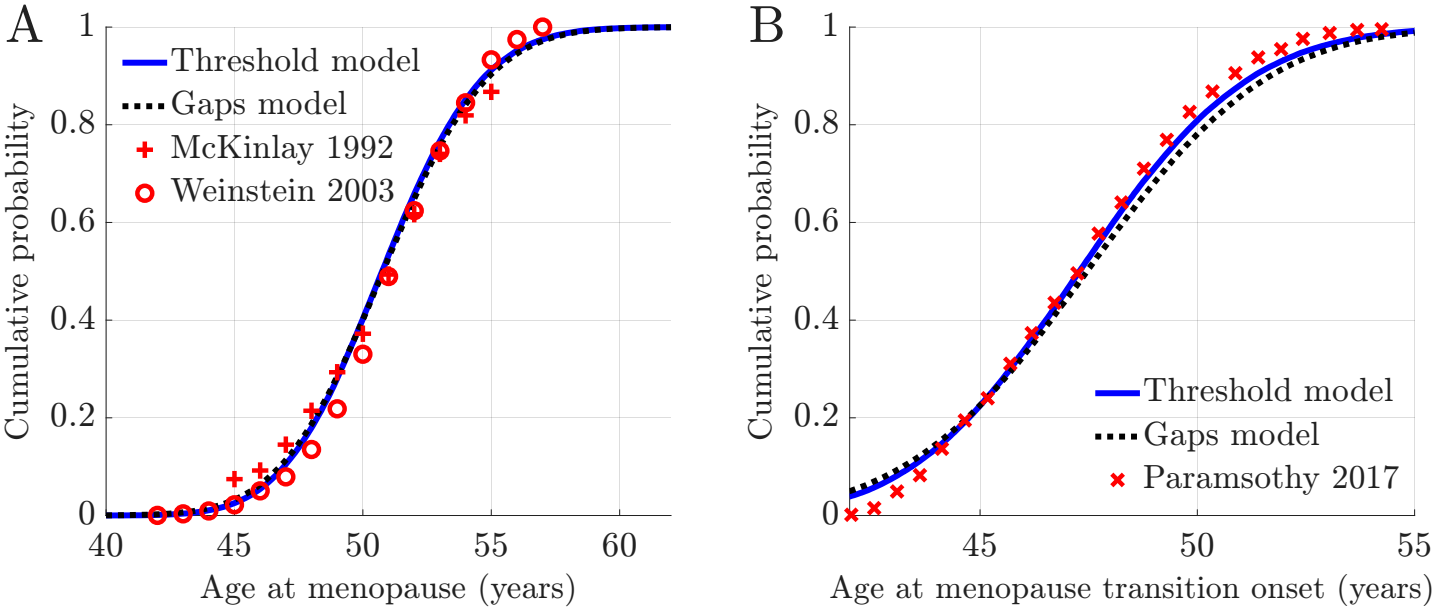
Simulated population distributions of ANM and AMT and empirical data [33, 35, 41]. In both panels, the solid blue curve is the threshold model, in which the milestone (menopause and menopause transition) occurs when the ovarian reserve (total across both ovaries) falls to *k* = 1000 follicles for the ANM (see Eq. (5)) or *k*_MT_ = 2600 follicles for the AMT (see Eq. (6)). In both panels, the dotted black curve is the gaps model, in which the milestone occurs the first time that *g*_ANM_ = 7.25 days (ANM) or *g*_MT_ = 3.75 days (AMT) elapse in which no follicle leaves the reserve of either ovary. Both models use the doubly exponential decay model in (1) with *N* and *R* independently drawn from Eqs. (2)-(3) with medians 6.5*×*10^5^ and 0.0396 year^*−*1^ and variability parameters *σ*_*N*_ = 0.5 and *σ*_*R*_ = 0.05, respectively. For the gaps model, both ovaries are followed together (starting supply *N*), and each curve is computed from 10^5^ simulated women. In both panels, the red markers are empirical distributions obtained from Refs. [33, 35, 41].

The solid blue curve in Figure 2B shows the distribution of the AMT when *N* and *R* vary according to the same log-normal distributions, with the same value of *σ*_*R*_ as in Figure 2A. The simulated AMT distribution resembles the empirically observed distribution (red markers) reported in Ref. [35]. Hence, the same model of ovarian reserve decay and population variability yields ANM and AMT distributions resembling those observed in actual populations.

The dotted black curves in Figure 2 show that the gaps model of ovarian aging milestones [30] also produces ANM and AMT population distributions resembling those observed in real women. The results in Figure 2A for the gaps model use the same distributions of *N* and *R* as above, but model ANM as the first time that there is a gap of 7.25 days in which no follicle leaves either ovary. The AMT for the gaps model in Figure 2B is modeled in the same way but with a gap of 3.75 days.

### 3.3 Halving the reserve modestly hastens menopause

The formula in (5) predicts that the ANM age *T* depends doubly logarithmically on the starting supply *N* . This very weak dependence means that large changes in starting supply produce comparatively small changes in ANM. For example, cutting the starting supply in half hastens menopause by only about 2.5 years,

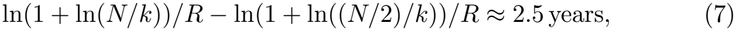

where (7) takes the median values for *N* and *R*. Hence, a 50% reduction in the ovarian reserve hastens ANM by a mere 5%. Similarly, increasing the starting supply from 2*×* 10^5^ (bottom 1% of population) to 2 *×*10^6^ (top 1% of population) increases the ANM by less than 8 years. That is, a 900% increase in starting supply produces a mere 17% increase in ANM.

Furthermore, this weak, doubly logarithmic dependence offers an explanation for the observation that a unilateral oophorectomy (UO) is associated with only a slightly earlier menopause age. Indeed, it was found that the average menopause age in women who had a UO is only 1.8 years earlier than women with two intact ovaries [38], compared with the hastening of 2.5 years in (7).

The doubly exponential model may also explain the empirically observed near independence between age at UO and age at menopause (see the very weak correlation in Figure 3 of Ref. [38]). To see this, observe that if menopause coincides with the first time that the ovarian reserve dips below the threshold of *k* follicles, then subtracting 50% of the reserve at birth versus subtracting 50% of the reserve at any later age yields the same menopause age, assuming merely that the subtraction happens when the reserve is larger than size 2*k*. Hence, the menopause hastening described in (7) indeed models the effect of a UO if the two ovaries have equal follicle numbers.

**Figure 3:**
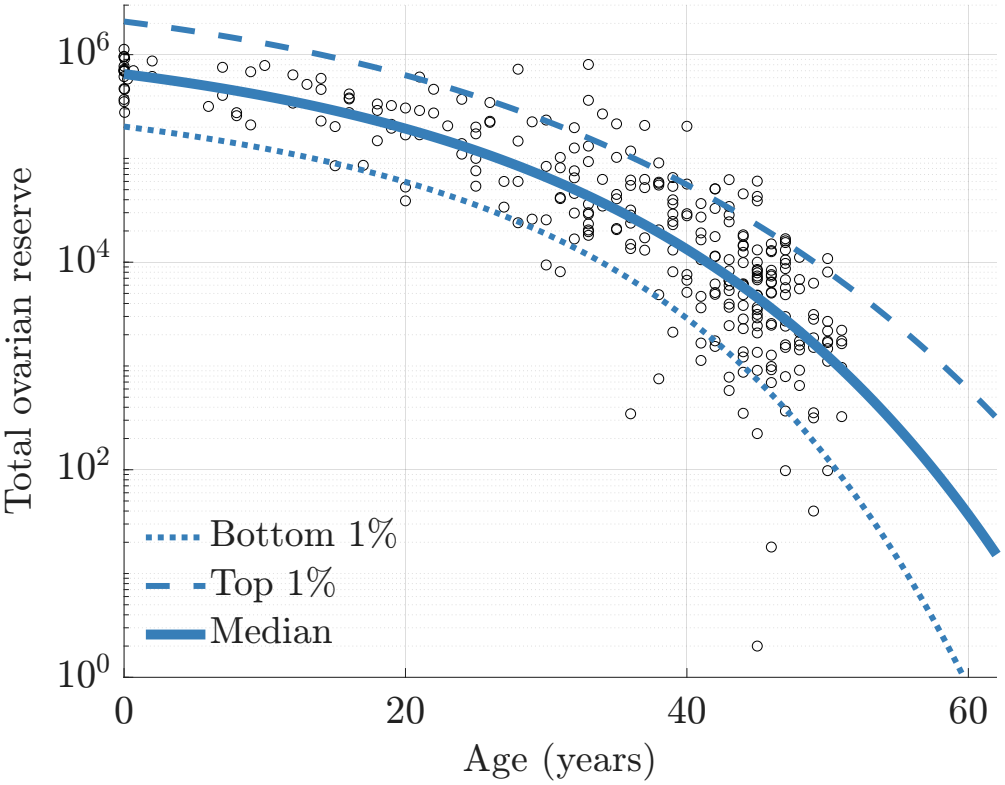
Doubly exponential decay model in Eq. (1) for the median (solid curve), top 1% (dashed curve), and bottom 1% (dotted curve) of a population in which the starting supply *N* and rate *R* vary according to the log-normal distributions in Eqs. (2)-(3), computed from 10^6^ simulated women. The circle markers are the histological follicle counts for 292 women shown in Figure 1. All counts and curves are totals per woman (both ovaries).

It is straightforward to generalize (7) to investigate the effects of a UO if the two ovaries do not have equal shares of the ovarian reserve. For example, if one third of all follicles are in one ovary and the other ovary contains the other two thirds of follicles, then the hastening becomes 4 years if the ovary with more follicles is removed (replace *N/*2 in (7) by *N/*3) and the hastening becomes 1.4 years if the ovary with fewer follicles is removed (replace *N/*2 in (7) by 2*N/*3). Note that this example of a one-third/two-thirds split of follicles between the two ovaries is more asymmetric than what is typically observed. Out of the 92 women whose ovaries were counted separately in Refs. [4, 5, 16], only 10 had divisions more asymmetric than one-third/two-thirds, and some of this asymmetry likely reflects counting error. Indeed, Hansen et al. found that the follicle count differences between a woman’s two ovaries were not significantly larger than the differences between repeated counts of the same ovary [18].

### 3.4 Variability in reserve size widens with age

The follicle count data in Figure 1 is highly variable. Moreover, the variability appears to increase over a woman’s lifetime, as the follicle counts between age-matched women vary roughly tenfold at birth and around a hundredfold near midlife.

Figure 3 shows that the doubly exponential model with log-normally distributed starting supply *N* and decay rate *R* qualitatively reproduces these two signatures of follicle count data. In particular, the 1st and 99th percentiles of the model distribution differ by a factor of approximately ten at birth, and this factor reaches one hundred by age 53. Hence, variation in *N* and *R* in the doubly exponential model captures some of the variability seen in follicle count data. However, of the 292 nonzero counts, 7.9% lie below the 1st-percentile model curve, and 8.9% lie above the 99th-percentile model curve, indicating that the model does not capture the full spread of the observed counts (see the Discussion section).

## 4 Discussion

In this paper, we developed and analyzed a new doubly exponential model of the decay of the ovarian reserve, and we assembled a new dataset of histological follicle counts after birth. This model involves a relatively simple mathematical formula that fits ovarian reserve data across postnatal life. Simultaneously, the model recapitulates population distributions for timing of menopause and the MT. This model also has the advantage of using fewer fitted parameters, reducing the likelihood of model “overfitting” where a model’s fit to available data is not generalizable to the process being studied.

The doubly exponential model also helps explain some curious features of ovarian aging. In particular, the model predicts that the ANM depends doubly logarithmically on the starting supply of oocytes, which means that large changes in starting supply produce only small changes in ANM. This offers an explanation for the comparatively mild population variability in ANM (coefficient of variation less than 10%) versus a large population variability in starting supply *N* (coefficient of variation more than 50%). Similarly, while it is intuitive that removing an entire ovary hastens menopause, it is a priori surprising that the empirically observed hastening associated with a UO is a mere 4% [38]. Indeed, “the relatively moderate decrease in age at menopause, however, is not understood” [38]. The doubly exponential model is compatible with this moderate decrease, as it predicts that halving the reserve only modestly has-tens menopause without any compensation by the remaining ovary. This does not establish that compensation by the remaining ovary is absent or unnecessary [24]. Note also that the model’s predicted hastening is reduced (respectively, increased) if the removed ovary had less (respectively, more) than half of the reserve. Related to the question of compensation, Grynberg et al. [17] found no altered ‘per-ovary’ and ‘per-follicle’ anti-Müllerian hormone production and antral follicle distribution in unilaterally oophorectomized women compared to matched controls. Last here, we showed that estimated variability in decay rate *R* between individuals is small, where most of the population is within 5% of the median value of *R*. This suggests that even in the absence of information about factors that might modify this rate in individuals, primordial follicle loss is a robust process that occurs within somewhat tight boundaries across the population.

Considering ovarian aging landmark events, doubly exponential reserve decay fits both historical “threshold” models and also our recent demonstration that “gaps” in the production of growing follicles from the ovarian reserve correspond to MT and ANM onset timing. In the threshold model [12], menopause is triggered when the ovarian reserve size falls below a critical number. This requires that the reproductive axis can sense the size of the remaining dormant reserve. In contrast, in the gaps model [30], menopause is triggered by a sufficiently long time interval elapsing without a new follicle leaving the reserve, with no explicit reference to the total follicle count. We favor the gaps model because of its physiological plausibility (e.g., loss of a regular supply of growing follicles required for reproductive cycling), and because it represents a mechanism by which the ovary can indeed “count” the number of remaining follicles. Since follicles exit the reserve at a rate roughly proportional to the reserve size, a sufficiently long gap is statistical evidence that the reserve has fallen below a critical size. The distinction between the threshold and gaps models mirrors two contrasting conceptions of the ovary, which we have elsewhere termed the Actively Regulated Ovary (ARO) and the Laissez-Faire Ovary (LFO) [29]. Under the ARO, a central organ-level controller orchestrates the decline of the reserve by monitoring the number of follicles in different stages of development [8]. Under the LFO, the decline of the reserve emerges from each follicle behaving stochastically and independently of other follicles [29].

Doubly exponential decay of the ovarian reserve as in the LFO can arise if each follicle exits at independent and identically distributed random times with a Gompertz probability distribution with scale *R* and unit shape. The Gompertz distribution includes an exponentially increasing probability of loss over time (its “hazard rate,” see below), and has been shown to result from the first passage time of certain diffusion processes modeling the first time that an accumulated “damage” crosses a threshold [23, 36]. Indeed, since ovarian reserve decay could be a mixture of loss due to growth and loss due to death, this increasing hazard rate could result from increasing cumulative damage. A prior model of ovarian aging similarly modeled follicle exit times as first passage times of diffusion processes modeling the first time that a stress response crosses a threshold [20].

It is illuminating to compare models of ovarian reserve decay in terms of their so-called “hazard rates.” The hazard rate describes the probability that a follicle that is still in the reserve will exit in the next instant of time (exit due to growth activation or follicle death via atresia). The hazard rate for the original biphasic model [12] abruptly increases at age 37.5 years (see the SI). For the doubly exponential model, the hazard rate is the exponentially increasing function of time *Re*^*Rt*^. Hence, the doubly exponential model can be understood as replacing the discontinuous hazard rate of the biphasic model by a smoothly increasing hazard rate. As a simple takeaway, the hazard rate *Re*^*Rt*^ means that the per-follicle hazard is approximately 4% per year at birth and doubles every (ln 2)*/R ≈*17.5 years. In particular, any follicle in the ovarian reserve at birth has an approximate 4% chance of exiting in the first year of postnatal life, and a follicle in the ovarian reserve at age 50 has an approximate 25% chance of exiting before age 51.

The follicle count data assembled here have important strengths. To our knowledge, they form the largest compilation of histological follicle counts after birth. Specifically, the data include 299 women from birth to age 55 years (7 of whom had estimated counts of zero), and each count is traced to a prior publication. Prior aggregations include 238 women in Ref. [7] (described as “the largest aggregation of human follicular data analyzed thus far” [7]) and at most 234 women in Ref. [40] (see SI Section S3.1). Furthermore, both ovaries were counted for 92 women in our dataset, which allowed us to confirm that doubling the count in a single ovary reliably estimates the total count across both ovaries.

The data and model presented here also have several limitations. For instance, the histological follicle counts underlying Figure 1 were collected across multiple studies using different counting protocols, follicle definitions, and tissue sampling strategies, as well as different sources (some data from autopsies and some from women having elective surgery) [4, 5, 14, 16, 18, 25, 37]. This heterogeneity plausibly contributes to the large scatter in the data, separate from genuine biological variability between women. Some of this scatter may stem from how different studies selected women. For instance, Gougeon et al. [16] required regular menstrual cycles with recent ovulation and Knowlton et al. [25] did not enroll women with more than 3 months of amenorrhea, and the counts in these two studies lie above our fitted curve on average. In contrast, 11 of the 17 women in Richardson et al. [37] were perimenopausal or postmenopausal (three of whom had zero follicles counted), and the nonzero counts in this study lie below our fitted curve on average. Nevertheless, the fitted rate constant and the median ANM change only slightly across reasonable choices of studies, follicle definitions, and treatment of the zero counts (see the SI). As a related point, about 17% of the 292 nonzero follicle counts lie outside the range between the bottom 1% and top 1% model curves in Figure 3, so there appears to be an additional source of variability in follicle counts not captured by the model. While this discrepancy might lead one to posit greater population variability in the follicle decay rate parameter *R* to better match the spread in follicle counts near midlife, doing so would yield simulated ANM and AMT distributions which are broader than those observed in populations of women. In addition, our lognormal choices for the population distributions of starting supply *N* and decay rate *R* were selected for simplicity and are not uniquely determined by the available data. Other distributions with comparable variability give similar results. Furthermore, for simplicity, we modeled *N* and *R* as independent across women. However, correlations between starting supply and decay rate could alter the simulated ANM and AMT distributions. As an additional limitation, our analysis did not model women with certain health conditions such as primary ovarian insufficiency (POI). However, it is plausible that women with POI could be described by the doubly exponential model with a large decay rate parameter *R* and/or a low starting supply *N*.

In addition to providing new information about the boundaries of follicle behavior and reproductive aging landmark timing, the doubly exponential model allows interrogation of physiological processes that can give rise to these boundaries. As shown and discussed, doubly exponential decay arises if individual primordial follicles exit the reserve at independent, random times, and this is consistent with stochastic threshold-crossing mechanisms of follicle loss. Determining how the independent, stochastic growth activation of human primordial follicles is controlled biologically is a high-priority next step.

## Authors’ roles

SDL and JJ participated in study design, execution, analysis, and manuscript drafting and review.

## Acknowledgments

SDL is supported by NSF DMS-2325258 and NSF DMS-2602237. JJ is supported by NSF DMS-2325259, NIH R21HD118658 and NIH R21HD120984. SDL and JJ are supported by the NIH Office of Research on Women’s Health (ORWH)/DPCPSI/Office of the NIH Director pursuant to Research Opportunity Announcement OTA-26-004, Computational Modeling of Hormone Homeostasis, award 1OT2OD042813. The views and conclusions contained in this document are those of the authors and should not be interpreted as representing official policies, either expressed or implied, of the NIH. During the preparation of this work, the authors used Claude (Anthropic) and Chat-GPT (OpenAI) to assist with compiling the follicle count dataset (transcribing counts from published tables, reading values from published figures, and tracing the values of a prior compilation to their sources), drafting and revising text, checking calculations, writing code, and formatting references. The mathematical model, analysis, interpretation, and conclusions are the authors’ own. All data, text, calculations, and code produced with this assistance were reviewed and verified by the authors, who take full responsibility for the content of this publication.

## Data availability

The follicle count dataset assembled for this study, giving for each woman the original source, follicle definition, and conversion to a total per woman, is provided as Supplementary Data and is archived at Zenodo (https://doi.org/10.5281/zenodo.24196625) [28]. All other data analyzed in this study were previously published and are available in the cited sources. No new histological data were generated. The analysis code is available from the corresponding author on reasonable request.

