## Supplementary Information for "Doubly exponential decay of the ovarian reserve and aging milestones"

### Supplemental Information for "Doubly exponential decay of the ovarian reserve and aging milestones"

Sean D. Lawley* Joshua Johnson†

November 21, 2026

#### S1 Extreme value theory

In the main text, we modeled the menopause age as the time when the ovarian reserve hit a critical threshold. More precisely, for a given model of ovarian reserve decay $F(t)$, the menopause age $T$ was defined as the time $T$ satisfying

$$F(T) = k. \tag{S1}$$

However, some prior models assume that each of the $N \gg 1$ follicles exits the reserve at independent and identically distributed (iid) random times, and the menopause age is defined to be the stochastic time $\mathcal{T}$ when the number of follicles remaining first falls below $k$ (see Ref. [17] for a review of such models). Since the physiologically relevant parameter regime is $1 \ll k \ll N$, it turns out that the stochastic time $\mathcal{T}$ has a probability distribution which is tightly centered on the deterministic time $T$ satisfying (S1). We now detail this argument.

Let $\{\tau_n\}_{n=1}^N$ be an iid sequence of realizations of a random variable $\tau$ with survival probability $S(t) = P(\tau > t) = F(t)/N$. For the doubly exponential model, $S(t) = \exp(1 - \exp(Rt))$, but we carry out our analysis for a general survival probability $S(t)$ that is differentiable with $f(t) = -S'(t) > 0$ for $t > 0$ with $S(0) = 1$ and $\lim_{t\to\infty} S(t) = 0$. Let $\mathcal{T}$ denote the $k$-th largest value of $\{\tau_n\}_{n=1}^N$ where $1 \ll k \ll N$. For any $t \geq 0$, define the number of realizations of $\tau$ which are larger than $t$,

$$M(t) = \sum_{n=1}^{N} 1_{\{\tau_n > t\}},$$

*Department of Mathematics, University of Utah, Salt Lake City, UT 84112 USA.

†Department of Obstetrics and Gynecology, University of Colorado School of Medicine, 12631 E 17th Avenue Building RC2, Room P15 3103 Aurora, Colorado 80045 USA.

where $1_{\{A\}}$ denotes the indicator function on the event $A$ (i.e. $1_{\{A\}} = 1$ if $A$ occurs and $1_{\{A\}} = 0$ otherwise). Notice that the $k$-th largest value $\mathcal{T}$ is larger than $t$ if and only if there are at least $k$ realizations of $\tau$ which are larger than $t$, i.e.

$$P(\mathcal{T} > t) = P(M(t) \geq k).$$

Therefore, we can estimate the distribution of $\mathcal{T}$ by estimating the distribution of $M(t)$. Notice that $M(t)$ is a Binomial random variable with $N \gg 1$ trials and trial success probability $S(t)$, and thus has mean $NS(t)$ and variance $NS(t)(1-S(t))$. Notice further that the $T$ defined by (S1) is given by $T = S^{-1}(k/N)$. Therefore, for $t$ near $T$, the count $M(t)$ has mean $NS(t) \approx NS(T) = k$ and variance $NS(t)(1-S(t)) \approx NS(T)(1-S(T)) = k(1-k/N) \approx k \gg 1$. Hence, for $t$ near $T$, the de Moivre-Laplace theorem implies that $M(t)$ is approximately normally distributed with mean $NS(t)$ and variance $k$, i.e.

$$P(\mathcal{T} > t) = P(M(t) \geq k) \approx \Phi\Big(\frac{NS(t) - k}{\sqrt{k}}\Big), \tag{S2}$$

where $\Phi(x) = \int_{-\infty}^{x} \frac{1}{\sqrt{2\pi}} e^{-z^2/2}\,\mathrm{d}z = \frac{1}{2}(1 + \mathrm{erf}(x/\sqrt{2}))$ is the cumulative distribution function of a standard normal random variable. Taylor expanding $S(t)$ for $t$ near $T$ gives

$$NS(t) \approx NS(T) + NS'(T)(t - T) = k - Nf(T)(t - T).$$

Plugging this Taylor expansion into (S2) gives

$$P(\mathcal{T} > t) \approx \Phi\Big(\frac{T - t}{\sqrt{k}/(Nf(T))}\Big),$$

which means that $\mathcal{T}$ is approximately normally distributed with mean $T = S^{-1}(k/N) = F^{-1}(k)$ and standard deviation $\sigma = \frac{\sqrt{k}}{Nf(T)}$, i.e.

$$\mathcal{T} \approx T + \sigma Z, \tag{S3}$$

where $Z$ is a standard normal random variable. Applying this argument to the doubly exponential model, we have $S(t) = \exp(1 - \exp(Rt))$ and thus $f(t) = -S'(t) = R\exp(1 + Rt - \exp(Rt))$ and $T = \ln(1 + \ln(N/k))/R$. Furthermore, taking $R = 0.0396\,\mathrm{year}^{-1}$, $k = 1000$ and $N = 6.5 \times 10^5$ as in the main text, we have that the mean of (S3) is $T = 50.8$ years and the standard deviation is $\sigma = 0.11\,\mathrm{year} = 1.3\,\mathrm{months}$.

To summarize, if one defines the menopause age as the stochastic time $\mathcal{T}$ which is the $k$th largest follicle exit time [17], then this stochastic time is centered at the deterministic time $T$ in (S1) and is within 0.3 years (about 3.5 months) of $T$ with probability 99%. Note that this calculation is for a fixed starting supply $N$ and rate parameter $R$; the variation in the population distribution of menopause age calculated in the main text is due to population variability in $N$ and $R$.

### S2 Reviewing prior models

We now review the eight prior models of the decay of the ovarian reserve that are compared with the doubly exponential model in Table S2. We also consider a more general doubly exponential model than the doubly exponential model considered in the main text.

#### S2.1 Biphasic model [8]

The biphasic model for the size of the ovarian reserve at age $t \geq 0$ is [8]

$$F(t) = \begin{cases} N \exp(-R_1 t) & \text{if } 0 \leq t < t_c, \\ N \exp(-R_1 t_c) \exp(-R_2(t - t_c)) & \text{if } t \geq t_c, \end{cases} \tag{S4}$$

where the four parameters are the rates $R_1 = 0.097/\text{year}$ and $R_2 = 0.237/\text{year}$, critical time $t_c = 37.5\,\text{years}$, and starting supply $N = 9.52 \times 10^5$. These four parameters were fit to the 110 follicle counts reported in Refs. [3,4,10,20]. The hazard rate of the biphasic model (the probability per unit time that a follicle that is still in the reserve exits) is the following piecewise constant function of time which abruptly increases at age $t_c = 37.5$ years,

$$-\frac{1}{F}\frac{\mathrm{d}F}{\mathrm{d}t} = \begin{cases} R_1 & \text{if } 0 \leq t < t_c, \\ R_2 & \text{if } t \geq t_c. \end{cases} \tag{S5}$$

For the doubly exponential model, the hazard rate is the following exponentially increasing function of time,

$$-\frac{1}{F}\frac{\mathrm{d}F}{\mathrm{d}t} = Re^{Rt}. \tag{S6}$$

Hence, the doubly exponential model can be understood as replacing the discontinuous hazard rate in Eq. (S5) by the smoothly increasing hazard rate in (S6).

#### S2.2 ODE model [7]

Faddy and Gosden later posited the following ordinary differential equation (ODE) model for reserve size $F(t)$ [7],

$$\frac{\mathrm{d}F(t)}{\mathrm{d}t} = -\Big(R_1 + \frac{R_2 - R_1}{1 + F(t)/F_c}\Big)F(t), \quad F(0) = N, \tag{S7}$$

where the four parameters are the rates $R_1 = 0.0595/\text{year}$ and $R_2 = 0.37495/\text{year}$, critical follicle number $F_c = 11780$, and starting supply $N = 7.0 \times 10^5$. These four parameters were fit to the 110 follicle counts reported in Refs. [3,4,10,20]. The ODE in (S7) is separable, and thus the age at which the reserve reaches a threshold $k$ has the following closed form

$$T_{\mathrm{ODE}}(k) = \frac{1}{R_2}\ln\Big(\frac{N}{k}\Big) + \Big(\frac{1}{R_1} - \frac{1}{R_2}\Big)\ln\Big(\frac{R_1 N + R_2 F_c}{R_1 k + R_2 F_c}\Big).$$

However, the reserve size $F(t)$ has no explicit formula and must be computed numerically.

#### S2.3 Compressed exponential model [12]

Hansen et al. [12] later posited the following compressed exponential model,

$$F(t) = N \exp(-(Rt)^p), \tag{S8}$$

where the three parameters are the power $p = 2.452$, rate $R = 0.04267$/year, and starting supply $N = 10^{5.717} = 5.2 \times 10^5$. These three parameters were fit to 122 follicle counts [12].

#### S2.4 Gaussian cumulative model [21]

Wallace and Kelsey [21] later proposed the following Gaussian cumulative decay curve to fit the follicle counts in their compilation,

$$F(t) = 2 \times 10^{f(t)},\ f(t) = \frac{a}{4}\left[1 + \operatorname{erf}\Big(\frac{t - b + c/2}{d_1\sqrt{2}}\Big)\right]\left[1 - \operatorname{erf}\Big(\frac{t - b - c/2}{d_2\sqrt{2}}\Big)\right], \tag{S9}$$

where the five parameters are $a = 5.56$, $b = 25.6$ years, $c = 52.7$ years, $d_1 = 0.074$ years, and $d_2 = 24.5$ years. In (S9), $\operatorname{erf}(z) = \frac{2}{\sqrt{\pi}}\int_0^z e^{-x^2}\,\mathrm{d}x$ denotes the Gauss error function. We note that formula (S9) corrects a typo in formulas (1) and (2) in Ref. [21] by putting a minus sign on the parameter $b$. We further note that the factor of 2 in Eq. (S9) converts the curve of Wallace and Kelsey, which describes one ovary, to a total per woman.

#### S2.5 Gompertz model [5]

In an important review of models of ovarian reserve decay [5], the following model was posited which the authors termed the Gompertz model,

$$F(t) = a(1 - \exp(-b\exp(-ct))),$$

where the three parameters $a = 302625.60$, $b = 3092.47$, and $c = 0.29\,\text{year}^{-1}$ were fit to follicle counts. Note that the starting supply for this model is $F(0) = a(1 - \exp(-b)) \approx a$ since $\exp(-b) \approx 10^{-1344} \approx 0$ for $b = 3092.47$.

#### S2.6 Random walk model [13]

The following "random walk model" of ovarian reserve decay was posited in Ref. [13],

$$\begin{aligned} F(t) &= \frac{N}{2}\left[1 + \operatorname{erf}\left(\frac{1 - Vt}{\sqrt{4Dt}}\right) - e^{V/D}\left(1 - \operatorname{erf}\left(\frac{1 + Vt}{\sqrt{4Dt}}\right)\right)\right], \\ &= \frac{N}{2}\left[\operatorname{erfc}\left(\frac{Vt - 1}{\sqrt{4Dt}}\right) - e^{V/D}\operatorname{erfc}\left(\frac{1 + Vt}{\sqrt{4Dt}}\right)\right], \end{aligned}$$

where $\mathrm{erf}(z) = \frac{2}{\sqrt{\pi}} \int_0^z e^{-x^2}\,\mathrm{d}x$ denotes the Gauss error function, $\mathrm{erfc}(z) = 1 - \mathrm{erf}(z)$, and the starting supply $N = 6.46 \times 10^5$ was taken to be the median of the population distribution near birth (the same population median quoted as $N = 6.5 \times 10^5$ in the main text), and the parameters $D = 0.004\,\mathrm{year}^{-1}$ and $V = 0.051\,\mathrm{year}^{-1}$ were fit to the postnatal follicle counts of Ref. [21] (these authors actually considered only a single ovary, which is equivalent to taking $N = 3.23 \times 10^5$).

#### S2.7 Chemical-kinetic model [18, 19]

The following "chemical-kinetic" model was posited in Ref. [19],

$$F(t) = Ne^{-\lambda t}(1 + \lambda t),$$

where the starting supply $N$ was as in the random walk model and the rate parameter $\lambda = 0.145\,\mathrm{year}^{-1}$ was fit to the postnatal follicle counts of Ref. [21]. This chemical-kinetic model was extended in Ref. [18], wherein the following model was posited,

$$F(t) = N \exp\Big(-\frac{\lambda}{a}(e^{at} - 1)\Big)\Big(1 - \frac{\lambda}{a}(1 - e^{at})\Big),$$

where the starting supply was as in the random walk model and the rate parameters $\lambda = 0.0906\,\mathrm{year}^{-1}$ and $a = 0.0217\,\mathrm{year}^{-1}$ were fit to the postnatal follicle counts of Ref. [21]. For a detailed review of the chemical-kinetic approach, see Ref. [17].

#### S2.8 More general doubly exponential model

The doubly exponential model in the main text is a special case of the following more general model,

$$F(t) = N \exp\big(\eta(1 - \exp(Rt))\big). \tag{S10}$$

That is, (S10) reduces to the doubly exponential model in the main text upon setting $\eta = 1$. In addition, when fitting the doubly exponential model in the main text to follicle counts, we fixed the starting supply at the previously reported population median $N = 6.5 \times 10^5$ [13] rather than letting it vary to improve the fit.

If we relax these two assumptions and fit (S10) to the 292 per-woman follicle counts with $R$, $\eta$, and $N$ all free, then the optimal values are

$$R_{\mathrm{opt}} = 0.0426\,\mathrm{year}^{-1}, \quad \eta_{\mathrm{opt}} = 0.85, \quad N_{\mathrm{opt}} = 6.5 \times 10^5.$$

Thus, the fitted starting supply agrees with the population median near birth, whereas $\eta$ moves somewhat below 1 and $R$ increases to compensate, so that the implied median ANM, $\ln(1 + \ln(N/k)/\eta)/R$ with $k = 1000$, is 50.6 years rather than 50.8 years. However, the fit improves only slightly (the SSE decreases from 84.08 to 83.96, or 0.1%), and $\mathrm{AIC_c}$ favors fixing $\eta = 1$ and $N = 6.5 \times 10^5$ (see Table S2).

### S3 Follicle count dataset

#### S3.1 The compilation of Wallace and Kelsey

| File S1 of Ref. [21] | | | Richardson et al. [20], Table 2 | | | | |
|---|---|---|---|---|---|---|---|
| Row | Age | Value | Subject | Age | One ovary | Twice one ovary | Diff. in $\log_{10}$ |
| 287 | 45 | 5,000 | A | 45 | 2,410 | 4,820 | +0.016 |
| 296 | 45 | 3,300 | B | 45 | 1,745 | 3,490 | −0.024 |
| 302 | 45 | 3,162 | E | 49 | 1,566 | 3,132 | +0.004 |
| 314 | 46 | 700 | I | 46 | 346 | 692 | +0.005 |
| 317 | 46 | 1,200 | H | 46 | 636 | 1,272 | −0.025 |
| 323 | 47 | 400 | C | 47 | 184 | 368 | +0.036 |
| 324 | 47 | 7,000 | D | 47 | 3,388 | 6,776 | +0.014 |
| 331 | 49 | 350 | K | 49 | 158 | 316 | +0.044 |
| 335 | 51 | 1,800 | F | 51 | 872 | 1,744 | +0.014 |

Table S1: The Richardson rows of File S1 of Ref. [21], which is labeled as counts per ovary, compared with the counts of Richardson et al. [20], who counted one ovary per woman. The last column is the $\log_{10}$ of the File S1 value minus the $\log_{10}$ of twice the one-ovary count. All nine values agree with twice the one-ovary count to within 0.05 in $\log_{10}$, whereas if the File S1 values were counts for one ovary, as labeled, these differences would be about $-\log_{10} 2 \approx -0.30$. Ages are in years; row 302 gives age 45 for subject E, aged 49. The File S1 values are round numbers (row 302 is $10^{3.5}$), which is consistent with reading them from the figure of Faddy et al. [8], which plots these women per pair of ovaries.

In an influential study, Wallace and Kelsey [21] compiled 336 values, including 288 at nonnegative ages and 11 values imposed at conception. Wallace and Kelsey made this compilation publicly available (File S1 of Ref. [21]), which made it possible to investigate the sources of these values. In doing so, we found some discrepancies. Such discrepancies are understandable in a large compilation because the original studies report counts in different ways (per ovary or per pair of ovaries, in tables or in figures, and for individual women or for age groups), and some values were available only from figures.

File S1 of Ref. [21] gives these 336 values in a table with 336 rows, but the study corresponding to each row is not stated. Nevertheless, Table 1 of Ref. [21] lists 52 values from Gougeon and Chainy [10] (ages 25 to 46 years, median 39.5) and 9 values from Richardson et al. [20] (ages 45 to 51 years, median 46), each described as one ovary. We assigned the rows that match the published tables of the other six studies in the compilation, and five further rows of Hansen et al. [12] (four rows with the value 300 at the ages of the four women with counts of zero, and one row (row 328) at the age of a woman whose count, halved, is 324 rather than the 342 entered). Apart from the 11 values imposed at conception, the remaining 61 rows divide into two groups with exactly the aforementioned numbers and age statistics. We call them the Gougeon rows (52 rows ranging from row 130 to row 321) and the Richardson rows (rows 287, 296, 302, 314, 317, 323, 324, 331, and 335). This reconstruction, with the matching criteria and the certainty of each assignment, is provided with our dataset (Supplementary Data) [16].

Below, rows are numbered from 1 to 336 as in File S1 of Ref. [21], and we describe the clearest examples of some discrepancies.

- **Totals for pairs of ovaries labeled as counts per ovary.** Richardson et al. [20] counted one randomly selected ovary per woman, and their Table 2 reports counts per ovary. The nine Richardson rows agree to within 0.05 in $\log_{10}$ with twice these counts, which are the values plotted per pair of ovaries by Faddy et al. [8], rather than with the counts themselves. We detail this in Table S1. Similarly, for the 32 Gougeon rows that we could match to individual women in Gougeon's data, the $\log_{10}$ difference between the File S1 value and that woman's total for both ovaries (from Table 1 of Gougeon et al. [11] where available, and otherwise from Figure 1 of Faddy [6]) has mean 0.00 and standard deviation 0.11, rather than the $-\log_{10} 2 \approx -0.30$ expected if these values were counts for single ovaries.

- **Possible duplicate women.** Some women of Block [3] appear to be represented both by their two single-ovary counts and by an additional total for both ovaries. For example, Block's case 27, aged 31 years, has primordial counts of 20,000 and 22,000 in her two ovaries (rows 165 and 169), and her total for both ovaries including small growing follicles is 44,690, which matches the value of 44,668 in row 171. Neither Table 1 of Gougeon et al. [11] nor Figure 1 of Faddy et al. [8] (as we read it) includes any woman of Gougeon aged 26 to 30 or 32 to 33 years, yet nine rows attributed to Gougeon have these ages, and eight of them closely match a woman of Block of the same age. Of the 20 Gougeon rows that we could not match to women of Gougeon, 13 rows closely and 4 approximately match totals for both ovaries of women of Block, and 3 are unresolved.

- **Zero counts.** The four zero counts of Hansen et al. [12] appear as 300 per ovary (rows 251, 264, 306, and 313).

Whatever the sources of the rows attributed to Gougeon, File S1 of Ref. [21] contains at most 234 eligible women at nonnegative ages: its 288 values there include two rows for each of the 43 women of Block [3] and the 6 newborns of Block [4] whose ovaries are entered separately, as well as 5 germ-cell records of Baker [1], and $288 - 43 - 6 - 5 = 234$. Under our candidate assignments, these 288 values correspond to 219 distinct records of our dataset (218 individuals and one group mean of Baker), 214 of which meet our eligibility criteria. Under the same assignments, our dataset includes 85 additional women: 52 of Knowlton et al. [14], 12 of Gougeon et al. [11] and 8 further women of Gougeon shown by Faddy et al. [8] that are not in File S1, 8 of Richardson et al. [20] (including the 3 with estimated counts of zero), and the 5 term newborns of Forabosco and Sforza [9], which File S1 places before birth.

Rather than modifying these values individually, we rebuilt a dataset from the original publications. These discrepancies change the fitted model only slightly. Indeed, fitting the compilation of Ref. [21] as published, after doubling it, gives $R = 0.0393\,\text{year}^{-1}$ rather than $0.0396\,\text{year}^{-1}$ (Table S3). We are grateful to Wallace and Kelsey for making their compilation publicly available, without which this comparison would not have been possible.

##### S3.2 Assembly of the dataset

The dataset [16] (Supplementary Data, with a description of every column) contains 331 records, including counts from preterm infants and germ-cell counts, each traced to the table or figure it came from. The counts are per woman (rather than per ovary) and were obtained as follows. When both ovaries were counted, we summed these two counts [3, 4, 11]. When one ovary was counted, we doubled it [9, 20], unless the authors had already done so [4, 12, 14]. The counts of Gougeon et al. [11] are the sums, over both ovaries, of non-growing follicles (their types B and B/C) and early-growing follicles (types C and D); these sums equal the counts of follicles smaller than 0.1 mm plotted by Faddy et al. [8]. Faddy et al. plotted 52 women of Gougeon (then unpublished); 43 of these were later tabulated by Gougeon et al. [11], and we read the other 9 from the figures of Faddy et al. [8] and Faddy [6]. The 52 women of Knowlton et al. [14] were read from their Figure 2. The digitized values were checked against values known from tables: readings from the figure of Faddy [6] agree with the tabulated counts of Block [3] and of Gougeon et al. [11] to within 0.005 on the $\log_{10}$ scale, and the digitized counts of Knowlton et al. reproduce the number of women, mean age, and mean count of every age group in their Table 1.

The fitting set used in the main text (292 women) contains every positive follicle count obtained at or after birth, regardless of menstrual status, except counts from five infants born preterm, whose ages measured from term are negative. It also excludes germ-cell counts and group means [1,2], and the seven women with estimated counts of zero. For each study, the follicle definition is the one used by Wallace and Kelsey [21] (and, for Knowlton et al. [14], that of Hansen et al. [12]).

#### S4 Ovarian reserve data fitting details and model comparison

As noted in the main text, the prior models were fit to different data sets than the doubly exponential model. We therefore refit each prior model to the same data set. The results are shown in Table S2. We now describe the fitting methods and results.

#### S4.1 Data

The fitting used the $n = 292$ per-woman follicle counts described in Section S3, at ages 0 to 51 years. Each count is the total across both ovaries.

#### S4.2 Fitting procedure

Let $\{(t_i, F_i)\}_{i=1}^{n}$ denote the ages and total reserve counts described above. For a model $F(t;\theta)$ with parameter vector $\theta$, we define the sum of squared errors on a logarithmic scale,

$$\mathrm{SSE}(\theta) = \sum_{i=1}^{n} \Big( \log_{10} F(t_i;\theta) - \log_{10} F_i \Big)^2 . \tag{S11}$$

Fitting on the logarithmic scale is appropriate here because the counts span several orders of magnitude. Indeed, an unweighted fit on the linear scale would be determined almost entirely by the counts near birth.

Parameters were estimated by minimizing (S11) with the Nelder–Mead simplex method [15] as implemented in the `fminsearch` routine of MATLAB R2026b (The MathWorks, Natick, MA, USA). To guard against local minima, each model was fit from $10^3$ starting points: the originally published parameter values, plus random perturbations obtained by multiplying each published value by $e^{\xi}$ with $\xi \sim \mathrm{Normal}(0, 0.35^2)$ drawn independently for each parameter.

Models with different numbers of parameters were compared using the corrected Akaike information criterion, denoted $\mathrm{AIC_c}$. Specifically, the $\mathrm{AIC_c}$ value for each model was computed according to

$$\mathrm{AIC} = n \ln\big(\mathrm{SSE}/n\big) + 2\kappa, \qquad \mathrm{AIC_c} = \mathrm{AIC} + \frac{2\kappa(\kappa+1)}{n-\kappa-1}, \tag{S12}$$

where $\kappa = p + 1$ and $p$ is the number of fitted parameters. The SSE for prior models was computed both at their originally published parameter values and after refitting, and $\mathrm{AIC_c}$ was computed from the refit SSE.

#### S4.3 Model comparison

Table S2 compares the doubly exponential model and prior models in terms of SSE and $\mathrm{AIC_c}$. The doubly exponential model in Eq. (1) of the main text has the smallest SSE of any model evaluated at its published parameter values. After refitting, the doubly exponential model has the lowest $\mathrm{AIC_c}$ of all models.

In particular, after refitting, the smallest SSE is 83.66 for the four-parameter ODE model [7], followed by 83.96 for the doubly exponential model with $\eta$ fitted and 83.97 and 83.98 for the four-parameter biphasic model [8] and the three-parameter Gaussian cumulative model [21], compared with 84.08 for the one-parameter doubly exponential model. Hence, no model improves on the doubly exponential fit by more than 0.5% in SSE. Finally, we point out that if $N$ is counted as a fitted parameter for the doubly exponential model, then its $AIC_c$ value increases by 2.04 (since $p$ increases from 1 to 2), but in this interpretation, $N$ should also be counted as a fitted parameter for the chemical-kinetic models [18, 19], which would increase the Mondal et al. [19] $AIC_c$ value by 2.04 (since $p$ increases from 1 to 2) and the Lyu et al. [18] $AIC_c$ value by 2.06 (since $p$ increases from 2 to 3).

| Model | $p$ | SSE (published) | SSE (refit) | $\Delta AIC_c$ |
|---|---|---|---|---|
| Doubly exponential, Eq. (1) | 1 | 84.08 | 84.08 | 0.00 |
| Doubly exponential, $\eta$ fitted | 2 | 84.08 | 83.96 | 1.63 |
| Doubly exponential, $N$ fitted | 2 | 84.08 | 84.01 | 1.81 |
| Chemical-kinetic, Lyu et al. [18] | 2 | 84.75 | 84.09 | 2.09 |
| Doubly exponential, $\eta$ and $N$ fitted | 3 | 84.08 | 83.96 | 3.68 |
| Gaussian cumulative [21] | $3^{\ddagger}$ | 84.85 | 83.98 | 3.77 |
| Compressed exponential [12] | 3 | 85.95 | 84.06 | 4.04 |
| ODE [7] | 4 | 85.63 | 83.66 | 4.70 |
| Biphasic [8] | 4 | 85.89 | 83.97 | 5.80 |
| Random walk [13] | 2 | 88.07 | 86.22 | 9.38 |
| Gompertz [5] | 3 | 107.17 | 86.06 | 10.91 |
| Chemical-kinetic, Mondal et al. [19] | 1 | 97.56 | 94.82 | 35.10 |

Table S2: Comparison of models of ovarian reserve decay against the 292 per-woman follicle counts (Section S3). Here $p$ is the number of fitted parameters, SSE is the sum of squared errors on a $\log_{10}$ scale defined in (S11), evaluated both at the originally published parameter values (for the doubly exponential model and its variants, $R = 0.0396\,\text{year}^{-1}$, $N = 6.5 \times 10^5$, and $\eta = 1$) and after refitting all parameters to these data, and $\Delta AIC_c$ is the difference in corrected Akaike information criterion (S12) relative to the best-scoring model, computed from the refit SSE. Rows are ordered by $\Delta AIC_c$. The rows labeled "$\eta$ fitted" and "$N$ fitted" are variants of the doubly exponential model in which the constraints $\eta = 1$ and $N = 6.5 \times 10^5$ are relaxed, as described in Section S2.8. $^{\ddagger}$We fitted all five parameters, but at the best fit, as at the published values, the factor $1+\text{erf}(\cdot)$ in Eq. (S9) equals 2 at every postnatal age, so only three combinations of the parameters ($a$, $b + c/2$, and $d_2$) are determined by postnatal counts, and we count $p = 3$. Counting $p = 5$ would give $\Delta AIC_c = 7.9$.

#### S4.4 The rate constant $R$

Minimizing (S11) over $R$ with $N = 6.5 \times 10^5$ held fixed gives

$$\widehat{R} = 0.039573\,\text{year}^{-1},$$

with SSE = 84.078. Resampling the 292 women with replacement ($10^4$ bootstrap samples) gives the 95% confidence interval 0.0388 to $0.0403\,\text{year}^{-1}$. In the main text we use this estimate rounded to three significant figures, $R = 0.0396\,\text{year}^{-1}$, which increases the SSE by less than 0.01%. With $k = 1000$ and $N = 6.5 \times 10^5$, this value gives a median ANM of 50.8 years.

#### S5 Sensitivity analyses

Table S3 shows the estimate $\widehat{R}$ of the rate constant (with $N = 6.5 \times 10^5$ fixed), the SSE, and the corresponding median ANM, $\ln(1 + \ln(N/k))/\widehat{R}$ with $k = 1000$, for alternative choices of data. The alternatives are: the compilation of Wallace and Kelsey [21] as published with every value doubled; the fitting set without particular studies or women; alternative follicle definitions; and median regression, which allows the seven women with estimated counts of zero to be included.

In median regression, the rate constant $R$ minimizes the sum of absolute deviations

$$\sum_i \left| \log_{10} F(t_i) - \log_{10} y_i \right|, \tag{S13}$$

instead of the sum of squared deviations, so that the fitted curve estimates the median count at each age rather than the mean of the $\log_{10}$ counts, in keeping with the interpretation of Eq. (1) as the median woman. Moving a count up or down without crossing the fitted curve changes the sum in (S13) by an amount that does not depend on $R$, and thus does not change the estimate (i.e. a count affects the estimate only through whether it lies above or below the fitted curve). The logarithm of a zero count is undefined, so we set each zero count to one follicle ($\log_{10} y_i = 0$), which lies below the fitted curve at the ages of all seven zero counts (42 to 55 years). Because $\log_{10} F(t_i)$ decreases as $R$ increases, the zero counts increase the estimate of $R$.

Across all of these alternatives, the median ANM ranges from 49.9 to 51.7 years. The fitted rate constant is most sensitive to the inclusion of the digitized counts of Knowlton et al. [14], which lie on average above the fitted curve, and to the use of median regression instead of least squares. Including the seven zero counts in the median regression increases $\widehat{R}$ by 1.2% and decreases the median ANM by 0.6 years.

| Data | $n$ | $\widehat{R}$ ($\text{year}^{-1}$) | SSE | ANM (years) |
|---|---|---|---|---|
| Compilation of Wallace and Kelsey [21] as published, doubled | 288 | 0.03927 | 67.33 | 51.2 |
| **Fitting set used in the main text** | **292** | **0.03957** | **84.08** | **50.8** |
| without the five women of Richardson et al. with fewer than 100 follicles | 287 | 0.03903 | 60.66 | 51.6 |
| without postmenopausal women | 291 | 0.03941 | 72.67 | 51.0 |
| without Knowlton et al. [14] | 240 | 0.04025 | 69.78 | 50.0 |
| without the 52 women of Gougeon [8, 11] | 240 | 0.03999 | 74.29 | 50.3 |
| without the 9 women of Gougeon not tabulated in [11] | 283 | 0.03963 | 81.07 | 50.8 |
| without all digitized counts | 231 | 0.04035 | 66.54 | 49.9 |
| Block [3]: primordial and growing follicles up to 0.1 mm | 292 | 0.03950 | 82.16 | 50.9 |
| Primordial or non-growing follicles only where available* | 292 | 0.03981 | 86.88 | 50.5 |
| Median regression, without the 7 zeros | 292 | 0.03888 | – | 51.7 |
| Median regression, with the 7 zeros | 299 | 0.03933 | – | 51.2 |

Table S3: Sensitivity of the fitted rate constant to the choice of data. The median ANM is $\ln(1+\ln(N/k))/\widehat{R}$ with $N = 6.5 \times 10^5$ and $k = 1000$. All rows except the last two are least-squares fits. *Non-growing follicles for Gougeon et al. [11] and primordial follicles for Richardson et al. [20] and Forabosco and Sforza [9]; the counts of Hansen et al. [12] and Knowlton et al. [14] include primary follicles and cannot be separated.

[11] A. Gougeon, R. Ecochard, and J. C. Thalabard. Age-related changes of the population of human ovarian follicles: increase in the disappearance

rate of non-growing and early-growing follicles in aging women. *Biology of Reproduction*, 50(3):653–663, 1994.
